# Vascular responses of hypercapnia challenge in mice

**DOI:** 10.1101/2024.09.20.614141

**Authors:** Xiuli Yang, Yuguo Li, Zhiliang Wei

## Abstract

Hypercapnia challenge with a few percent of carbon dioxide is popularly used in physiological studies to investigate the dynamic vascular properties. A typical hypercapnia experiment can be divided into four phases: baseline (regular air), transition (CO_2_ enriched), hypercapnia (CO_2_ enriched), and recovery (regular air). Cerebrovascular reactivity (CVR), denoting the percentage of functional changes between baseline and hypercapnia phases, can be measured to assess vascular health. In this study, we focus on the transition and recovery phases to track the built-up of cerebral blood flow (CBF) increase induced by CO_2_-enriched gas and recovery of CBF after returning to regular air. Dynamic features were compared with those of another potent vasodilatory agent, acetazolamide. Our results reveal that 5 min is sufficiently long to ensure 95% built-up of CBF increase under 5% CO_2_, but it takes much longer to recover to baseline.

## 1. Introduction

Magnetic resonance (MR) technique is a versatile tool for various applications,^1-5^ among which MR measurements provide a rich resource of biomarkers for denoting pathogeneses.^6-9^ Structural markers represent the early applications of MR imaging (MRI) based on its non-invasive nature.^4,10^ Due to its special mechanism to generate resonance signals, MRI has excellent imaging depth for *in-vivo* observations, making it a suitable tool for investigating the pathologic progression of diseases associated with changes in deep brain regions.^11-14^ With the technical developments in the past few decades, functional imaging markers targeting at a specific physiological aspect attract broad attentions^15,16^ and start to play important roles in the clinical and mechanistic studies, such as glucose transportation and oxygen metabolism in Alzheimer’s disease,^8,17,18^ blood-brain barrier permeability in cancer,^19,20^ and cerebrovascular reactivity (CVR) in vascular cognitive impairment and dementia.^21,22^

Gas with enriched CO_2_ content serves as a broadly utilized stimulant to examine CVR, termed as hypercapnia challenge.^23^ Specifically, the combination of MR recording, such as blood-oxygenation-level-dependent (BOLD)^24,25^ and pseudo-continuous arterial spin labeling (pCASL)^26,27^, with hypercapnia challenge provides a quantitative measurement for CVR.^23^ CVR denotes the percentage of functional change per unit stimulant and is presented with different units subject to the MR reading and vasoactive intervention. For example, CVR has a unit of %BOLD/mmHg when BOLD and capnograph are used.^23^ If a vasodilatory agent, such as acetazolamide (ACZ), is used as an alternative to the hypercapnia challenge, unit of CVR measurement becomes %/[µg/ml].^28^ Hypercapnia is often preferred for CVR measurements due to the simple implementation. Typically, MR recordings are collected before and after the hypercapnia challenge, with an appropriately chosen delay time in between.^29^ In literature, blood flow changes during this delay is understudied. Therefore, in this study, we aim to characterize the built-up of cerebral blood flow (CBF) increase induced by the hypercapnia challenge and the recovery of CBF after returning to normocapnia.

## 2. Methods

### 2.1. General procedures

The experimental protocols involved in this study were approved by the Johns Hopkins Medical Institution Animal Care and Use Committee and conducted in accordance with the National Institutes of Health guidelines for the care and use of laboratory animals. Data reporting followed the ARRIVE 2.0 guidelines. All mice had free access to food and water and were housed in a quiet environment with a 12-h day/night cycle. A total number of 11 C57BL/6 mice (5 females and 6 males; age: 4-7 months; body weight: 21-28 g) were used. Hypercapnia experiments can be divided into four different phases: baseline (with regular air), transition (with CO_2_-enriched gas), hypercapnia (with CO_2_-enriched gas), and recovery (returning to regular air) (Figure 1A).

**Figure 1.**
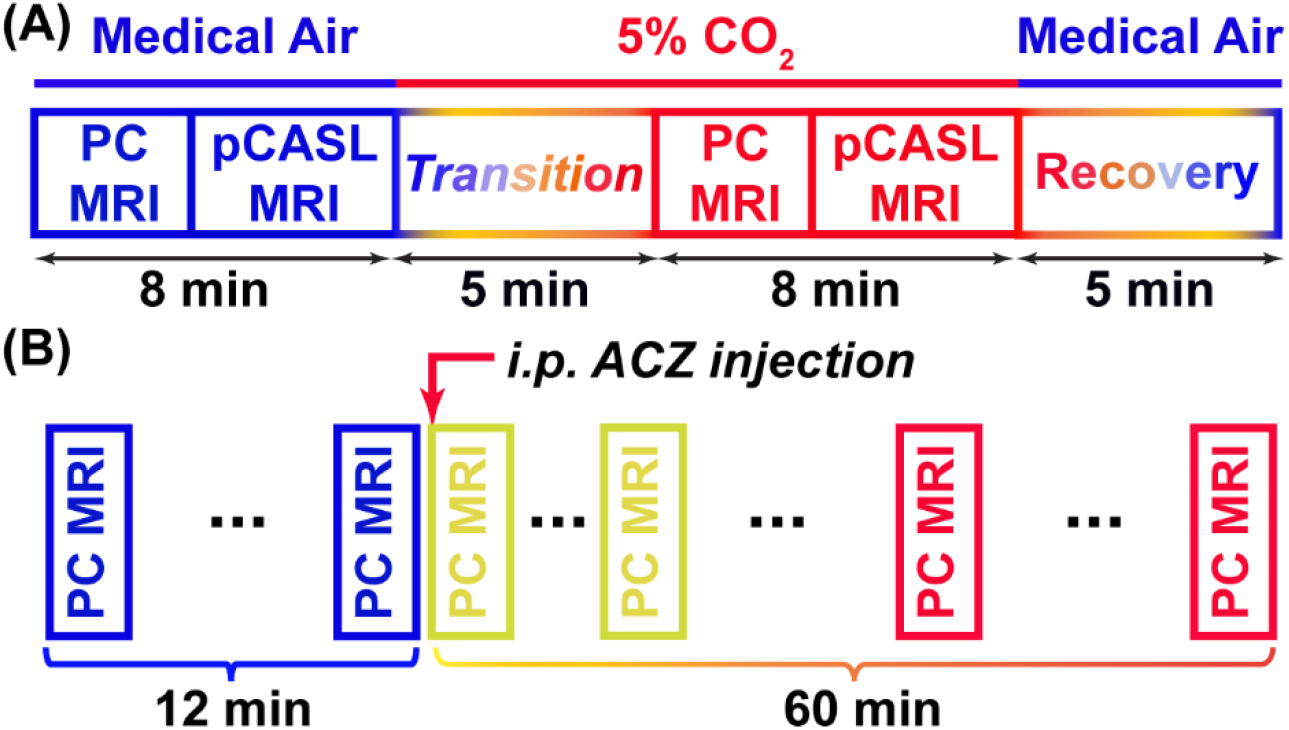
Schematic diagrams of study designs. PC: phase contrast; pCASL: pseudo-continuous arterial spin labeling.

### 2.2. MRI experiments

All MRI experiments were performed on an 11.7T Bruker Biospec system (Bruker, Ettlingen, Germany) with a horizontal bore as well as an actively shielded pulse field gradient (maximum intensity of 0.74 T/m). Images were acquired using a 72-mm quadrature volume resonator as the transmitter, and a four-element (2×2) phased-array coil as the receiver. The B_0_ homogeneity over the mouse brain was optimized with a global shimming (up to 2^nd^ order) based on a pre-acquired subject-specific field map. For mice scanned in the same day, randomization was performed to determine the experimental order following a reported scheme.^30^ Inhalational isoflurane was used throughout MRI scans.1.5-2.0% isoflurane was used as an induction for 15 min. At the 10^th^ minute under induction, the mouse was placed onto a water-heated animal bed with temperature control and immobilized with a bite bar, ear pins, and a custom-built 3D-printed holder before entering the magnet. After the anesthetic induction, the isoflurane level was adjusted within 1.0-1.3% to maintain respiration rates below 130 breath/min. Isoflurane was delivered with medical air (21% oxygen and 78% nitrogen) at the flow rate of 0.75 L/min. Respiration rate was observed during the experiment by a MR-compatible monitoring & gating system (SA Instruments, Stony Brook, USA).

CBF was evaluated with phase-contrast (PC) MRI covering the three major feeding arteries, i.e., left internal carotid artery (LICA), right internal carotid artery (RICA), and basilar artery (BA), in separate scans to collect corresponding through-plane velocity maps.^31^ Prior to the PC scans, we first performed a coronal time-of-flight (TOF) MRI angiogram (9 slices, slice thickness = 0.5 mm, no inter-slice gap, TR/TE = 45/2.6 ms, scan duration = 2.6 min) to visualize the feeding arteries. Then, a sagittal TOF (single slice, tilted to contain the targeted artery identified from coronal TOF images, thickness = 0.5 mm, TR/TE = 60/2.5 ms, scan duration = 0.4 min) was applied to visualize the in-plane trajectory of the targeted artery. Based on the reference TOF images (coronal and sagittal), PC MRI was then positioned and performed using the parameters of: TR/TE = 15/3.2 ms, FOV = 15×15 mm^2^, matrix size = 300×300, slice thickness = 0.5 mm, number of averages = 4, dummy scan = 8, receiver bandwidth = 100 kHz, flip angle = 25º, partial Fourier acquisition factor = 0.7, and scan duration = 0.6 min per artery.^32,33^ Number of averages might be changed to provide other temporal resolutions subject to the study purpose later.

Brain volume was estimated using a T_2_-weighted fast spin-echo MRI protocol: TR/TE = 3000/11.0 ms, FOV = 15×15 mm^2^, matrix size = 128×128, slice thickness = 0.5 mm (without inter-slice gap), echo spacing = 5.5 ms (4 spin echoes per scan), 35 axial slices, and scan duration = 1.6 min.^34^

Regional perfusion was evaluated by a two-scan pseudo-continuous arterial spin labeling (pCASL) MRI,^35^ which was optimized to minimize the influence of magnetic field inhomogeneity. First, a pre-scan was performed to optimize the phases of labeling pulses in the control and labeled scans. Then, the scan focusing on regional perfusion was performed with the following parameters: TR/TE = 3000/11.8 ms, labeling duration = 1800 ms, FOV = 15×15 mm^2^, matrix size = 96×96, slice thickness = 0.75 mm, labeling-pulse width = 0.4 ms, inter-labeling-pulse delay = 0.8 ms, flip angle of labeling pulse = 40°, post-labeling delay = 300 ms, two-segment spin-echo echo-planar-imaging acquisition, partial Fourier acquisition factor = 0.7, number of average =25, and scan duration = 5.0 min.^36^

### 2.3. Experimental design

We aimed to characterize the dynamic changes of CBF during the transition and recovery phases (N=10; 5 females and 5 males). Detailed design was outlined in Figure 1A. PC (covering three arteries) and pCASL scans were performed to measure baseline CBF under the medical air. Subsequently, gas containing 5% CO_2_ was administered. A 5-min delay was allocated for the vasoactive effect of hypercapnia to reach steady state, denoted as transition phase. PC and pCASL scans were repeated after the transition phase. Finally, gas was switched back to medical air and a 5-min delay (denoted as recovery phase) was utilized. During the transition and recovery phases, a single-artery PC focusing on LICA was repeatedly scanned (16 repetitions with a temporal resolution of 0.3 min). Additionally, a proof-of-principle checking on the dynamic CBF change after i.p. ACZ injection was implemented as Figure 1B. Repeated single-artery PC MRI focusing on LICA was implemented at a temporal resolution of 2.4 min with 30 repetitions, and the i.p. injection was administered at the starting of the 6^th^ repetition.

### 2.4. Data processing

Data processing of PC MRI utilized a custom-written graphic-user-interface (GUI) tool developed in MATLAB following the established procedures.^37^ The artery of interest was first delineated manually on a complex-difference image, which provided clear contrast between the vessel and surrounding tissue. This mask was then applied to the phase velocity map, and integration of arterial voxels yielded blood flow through the targeted artery in ml/min. Summing the blood flow values from the three major feeding arteries provided the total blood flow to brain. To normalize the brain-size differences and obtain unit-mass CBF values, the total blood flow was divided by the brain weight, which was calculated as the product of brain volume and brain tissue density (1.04 g/ml^38^). The global CBF value was reported in the unit of milliliters per 100 g brain tissue per minute (ml/100g/min).

Processing of pCASL data followed the reported procedures.^37^ Briefly, pair-wise subtraction between control and labeled images (i.e., *M_ctr_ − M_lbl_*) were first applied to yield a difference image, which was then divided by a M_0_ image (obtained by scaling the control image) to provide a perfusion index image, i.e., 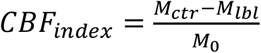. The perfusion index maps were co-registered and normalized to a mouse brain template^39^ and then resized to recover original acquisition resolutions. The normalized perfusion index maps were rescaled by reference to the global CBF (from PC MRI) to obtain absolute-value CBF, specifically, 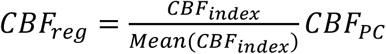. Regional-of-interest (ROI) was drawn on the averaged control images to encompass isocortex, hippocampus, thalamus, hypothalamus, and striatum by reference to the mouse brain atlas (https://atlas.brain-map.org/). Voxel-wise CBF values within each ROI were averaged to estimate the corresponding perfusion levels.

## 3. Results

### 3.1. Built-up of CBF increase after hypercapnia

Figure 2 illustrates the dynamic CBF increases associated with hypercapnia. Velocity maps (Figure 2A) exhibit prominent increases in flowing velocities. Figure 2B presents the percentage change of CBF during the transition phase. The dynamic data were fitted into the function of 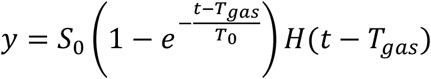, where S_0_ denotes the vasodilation amplitude, *T*_gas_ the arrival time of vasoactive effect, and *T*_0_ the built-up time. According to the fitting, vasoactive effect occurs at 1.35 min, which is attributed to the gas travelling from the gas tank to the mouse lung along the gas tubing and the gas exchange in the pulmonary alveoli. After *T*_gas_, CO_2_ molecules are dissolved in blood to modulate pH and ion-channel activities, thus inducing vasodilation.^40^ The built-up time *T*_0_ is 1.22 min, indicating the time required to reach 63.2% of the maximal CBF change. The percentage of CBF change at the end of a 5-min transition period has reached 95% of the maximal CBF change. Therefore, 5-min transition is recommended when running hypercapnia experiments with 5% CO_2_ in mice.

**Figure 2.**
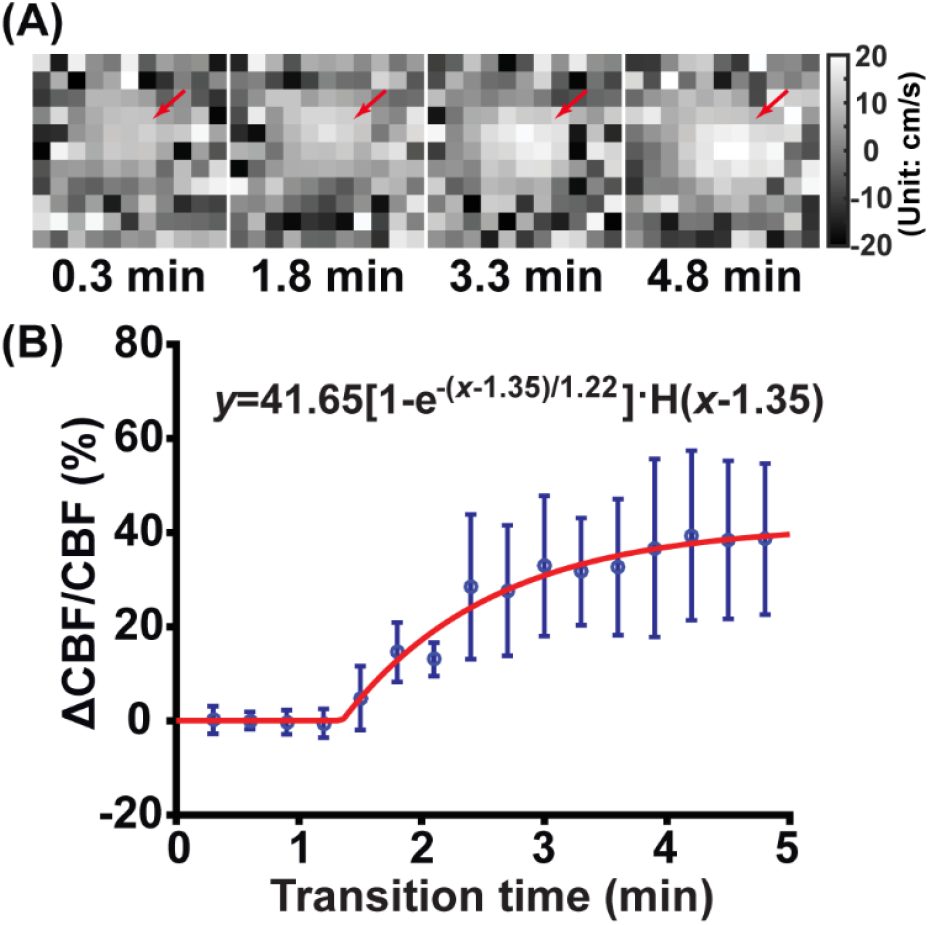
Characterization of dynamic CBF changes in the transition phase (N=10). (A) displays the velocity maps at 0.3, 1.8, 3.3, and 4.8 min. (B) percentage change of CBF as a function of transition time. Error bars represent standard error over mice (N=10). H(x) denotes the Heaviside step function.

**Figure 3.**
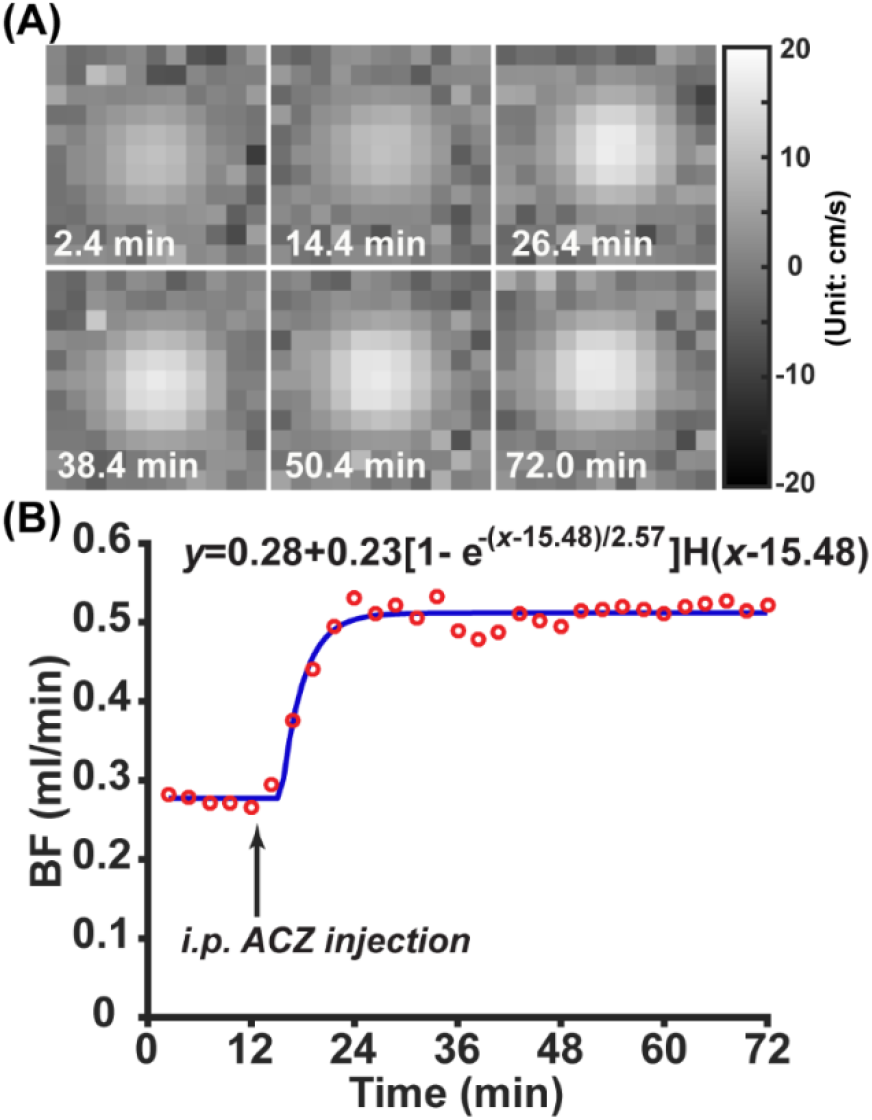
Dynamic CBF change with i.p. ACZ injection. (A) displays the velocity maps at 2.4, 14.4, 26.4, 38.4, 50.4, and 72.0 min. The i.p. ACZ injection was executed at 12^th^ min. (B) blood flow (BF) changes as a function of time. Red dots represent original BF values and blue line represent the fitted line. H(x) denotes the Heaviside step function.

Under the i.p. ACZ injection, *T*_ACZ_ (counterpart of *T*_gas_ in hypercapnia experiment) is 15.48-12.00=3.48 min, denoting the time it takes for ACZ to travel from enterocoelia to capillaries. In contrast, *T*_ACZ_ become shorter when intravenous (i.v.) ACZ injection was used. It seems that direct gas inhalation is associated with a swifter vascular response than the i.p. ACZ injection. The *T*_0_ associated with i.p. ACZ injection (2.57 min) is longer than the i.v. ACZ injection (1.62 min^28^). This is primarily because ACZ appears instantly in vasculature with i.v. injection but gradually accumulate in the case of i.p. injection, leading to different plasma levels of ACZ. On the other hand, *T*_0_ associated with hypercapnia is shorter than that with i.v. ACZ injection, possibly suggesting that altering blood pH by inhibiting carbonic anhydrase is a slower process than exchanging CO_2_ into blood via lung alveolar with a high partial pressure. A 5-min transition period will reach 41% and 95% of the maximal CBF change for i.p. and i.v. ACZ injections, respectively. When the i.p. injection is employed, the minimal transition time required to reach 95% effect is 11.2 min.

### 3.2. Recovery of CBF after switching to normocapnia from hypercapnia

Figure 4 illustrates the dynamic CBF decreases after cutting off hypercapnia. Velocity maps (Figure 4A) exhibit slight decreases in the flowing velocities. Figure 4B presents the percentage change of CBF during the recovery phase. The dynamic data were fitted into the function of 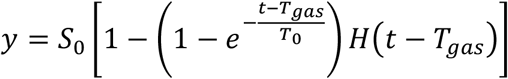, where *S*_0_ denotes percentage of CBF change at the initial state, *T*_gas_ the retarding time of vasoactive effect, and *T*_0_ the fade-away time. Since the recovery phase is the opposite process of the transition phase, the retarding time is assumed to be the same with arrival time (1.35 min). Data fitting reveals an averaged *T*_0_ of 7.27 min, which is 5.96 folds of the *T*_0_ in transition phase, suggesting that the removal of CO_2_ from blood after discontinuing 5% CO_2_ is a prominently slower process than the dissolution of CO_2_ into blood in lung alveolar. At the end of the 5-min recovery period, the averaged percentage CBF change is 22.8%. In contrast, percentage CBF change due to natural anesthetic accumulation at the same time point is 7.50%. Such a large discrepancy supports the notion that the high percentage CBF change at the end of a 5-min recovery period is due to the slow diminishment of hypercapnia effect instead of the anesthetic accumulation effect.

**Figure 4.**
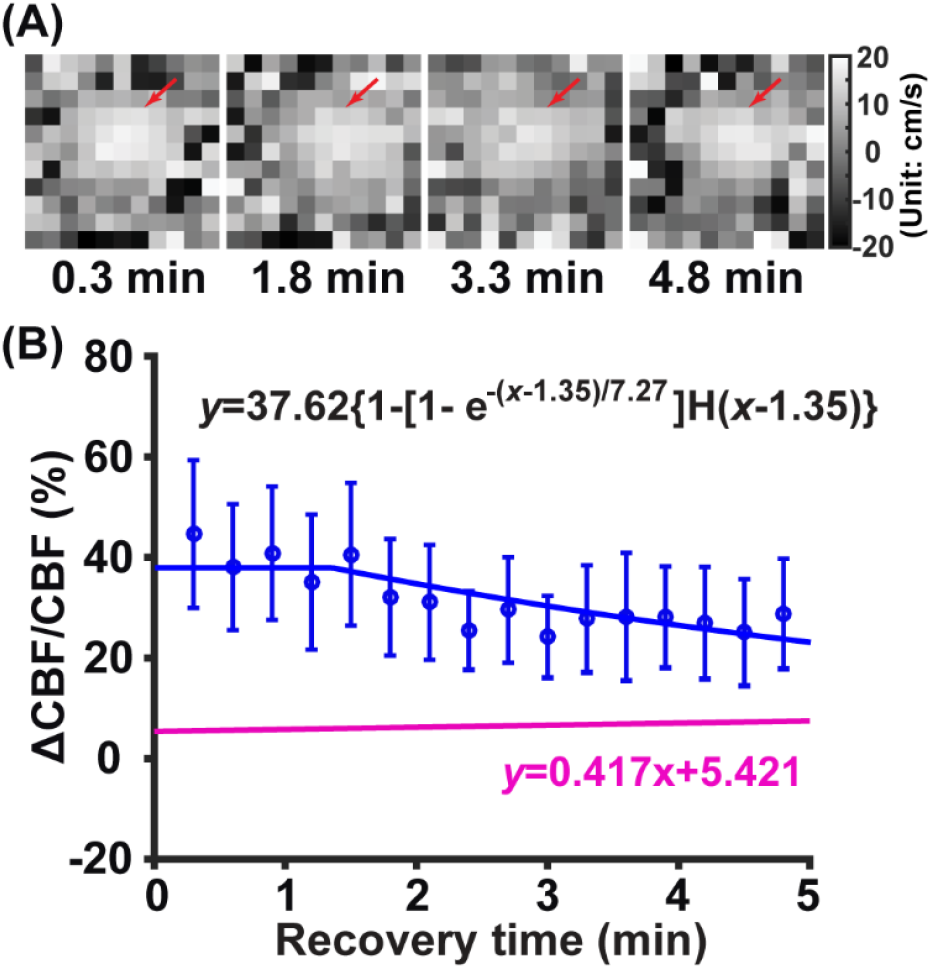
Characterization of dynamic changes in the recovery phase (N=10). (A) displays the velocity maps at 0.3, 1.8, 3.3, and 4.8 min. (B) percentage change of CBF as a function of recovery time. Error bars represent standard error over mice (N=10). Magenta line represents the fitted line of percentage CBF change due to natural anesthetic accumulation (without hypercapnia or ACZ injection). H(x) denotes the Heaviside step function.

### 3.3. Perfusion changes pre- and post-transition

Figure 5 presents the comparison between CBF maps pre- and post-transition. Consistent increases in perfusion can be observed across various brain regions, particularly in the cortical region. Such perfusion changing patterns resemble those following the i.v. ACZ injection.^28^

**Figure 5.**
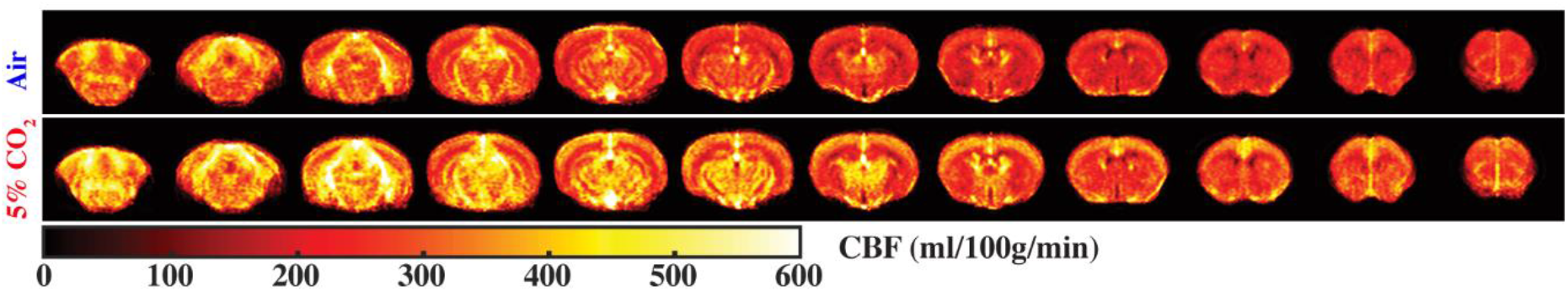
Averaged CBF maps pre- and post-transition (N=10).

## Discussion

We have investigated the transition and recovery phases of hypercapnia with quantitative MRI. It has been found that the vasodilatory effect of hypercapnia challenge has a swift built-up along with a slow diminishment.

Different facility may install gas tubing in different lengths, which will affect the arrival time of vasoactive effect. Assuming a tubing system with a homogeneous 5 mm internal diameter, gas can be delivered over 38.20 m in 1 min at the flow rate of 0.75 L/min. Considering that the gas room and scanner room are often located adjacently for experimental convenience, a gas system with 38.20 m should be sufficiently long for most facilities. Therefore, adding 1 min to the 5-min transition should be able to fit for the practical experimental setup of different facilities (namely, 6-min transition was recommended for other facilities as a general suggestion).

Mechanisms to induce vasodilation underlying hypercapnia challenge and ACZ injection are similar except for the initial steps to alter the blood pH environment. Under the hypercapnia challenge, due to the higher partial pressure of CO_2_ in the lung alveolar, more CO_2_ will be dissolved in blood and then exist in the form of carbonic acid.^41,42^ In the case of ACZ injection, the partial pressure of CO_2_ in the lung alveolar is not affected by the ACZ. Instead of dissolving more CO_2_ into the blood, the normal release of CO_2_ from blood to lung alveolar by decomposing carbonic acid is significantly perturbed because ACZ is a carbonic anhydrase inhibitor.^43,44^ From these comparisons, it can be concluded that hypercapnia challenge and ACZ injection lead equivalent modulation to the blood microenvironment by either introducing CO_2_ dissolution or blocking CO_2_ clearance.

Due to the slow diminishment of hypercapnia effect, a lengthy recovery phase is necessary for repeated CVR measurements. With the presence of anesthetic accumulation effect,^28^ CBF will not return to its value at the baseline phase. In literature, the half-life of ACZ in mice (in terms of the vasodilatory effect) is approximately 55 min.^28^ Therefore, it seems that CVR measurements using hypercapnia challenges or ACZ injections are better implemented at the end of experimental sessions alongside other sequences to minimize potential confounding.

Physiological MRI is an active subfield attracting more and more research interest because physiological measurement are sensitive and promising markers to track various pathologies or processes.^34,37,45-50^ Centering the microcirculation, OEF, CBF, CVR, and BBB permeability constructs a multiparametric insight into the complete landscape of microvascular function. These measurements can be achieved with non-invasive and non-contrast MRI techniques, thus avoiding the daunting task of inter-modality co-registration. Additionally, the relevant MRI techniques are fully translatable to clinic studies due to the parallel efforts of methodological development and optimization in both human and animals.^36,51-53^

In recent years, advanced signal processing algorithms including artificial intelligence has been broadly used in MRI to improve the technical performance.^54-57^ CVR mapping will also benefit from these state-of-the-art processing algorithms due to the general tradeoff among scan duration, spectral resolution, and signal-to-noise ratio. An improvement in either of these can translate into the others subject to better fulfill the study purposes.

## Conclusion

A transition period of 5 min is recommended for fully inducing the vasodilatory effect of hypercapnia. Hypercapnia challenge is associated with a swift built-up (63.2% increase in 1.22 min) and a slow diminishment (63.2% reduction in 7.27 min).

